# Experience sampling reveals the role that covert goal states play in task-relevant behavior

**DOI:** 10.1101/2023.07.12.548563

**Authors:** Brontë Mckeown, Will H. Strawson, Meichao Zhang, Adam Turnbull, Delali Konu, Theodoros Karapanagiotidis, Hao-Ting Wang, Robert Leech, Ting Xu, Samyogita Hardikar, Boris Bernhardt, Daniel Margulies, Elizabeth Jefferies, Jeffrey Wammes, Jonathan Smallwood

**Affiliations:** Psychology Department, Queen’s University, Kingston, K7L 3N6, Canada; Neuroscience, Brighton and Sussex Medical School, University of Sussex, Brighton, BN1 9RH, UK; CAS Key Laboratory of Behavioural Science, Institute of Psychology, Chinese Academy of Sciences, Beijing, 100101, China; Department of Psychiatry and Behavioral Sciences, Stanford University, California, 94305, USA; Department of Psychology, Durham University, Durham, DH1 3LE, UK; School of Psychology, University of Sussex, Brighton, BN1 9QH, UK; Centre de recherche de l’institut Universitaire de gériatrie de Montréal (CRIUGM), Montréal, Québec, H3W 1W5, Canada; Centre for Neuroimaging Science, King’s College London, SE5 8AF, UK; Center for the Developing Brain, Child Mind Institute, New York, 10022, USA; Department of Neurology, Max Planck Institute for Human Cognitive and Brain Sciences, Leipzig, 04103, Germany; Montreal Neurological Institute, McGill University, Montreal, H3A 2B4, Canada; Integrative Neuroscience and Cognition Center (UMR 8002), Centre National de la Recherche Scientifique (CNRS) and Université de Paris, Paris, 75006, France; Department of Psychology, University of York, York, YO10 4LX, UK

**Author notes:** **Corresponding author:** Brontë Mckeown.

## Abstract

Cognitive neuroscience has gained insight into covert states using experience sampling. Traditionally, this approach has focused on off-task states, however, task-relevant states are also maintained via covert processes. Our study examined whether experience sampling can also provide insights into covert goal-relevant states that support task performance. To address this question, we developed a neural state-space, using dimensions of brain function variation, that allows neural correlates of overt and covert states to be examined in a common analytic space. We use this to describe brain activity during task performance, its relation to covert states identified via experience sampling, and links between individual variation in overt and covert states and task performance. Our study established activity patterns within association cortex emphasizing the fronto-parietal network both during target detection and a covert state of deliberate task focus which was associated with better task performance. In contrast, periods of vigilance and a covert off-task state were both linked to activity patterns emphasizing the default mode network. Our study shows experience sampling can not only describe covert states that are unrelated to the task at hand, but can also be used to highlight the role fronto-parietal regions play in the maintenance of covert task-relevant states.

## Introduction

Contemporary cognitive neuroscience has established the neural correlates of hidden states such as mind-wandering ^1^, or more recently, mind-blanking ^2^, by associating brain activity with patterns of thought described by experience sampling ^3-5^. These states are a natural target for experience sampling studies because they help establish the correlates of states that occur spontaneously, and so are hard to evoke using standard experimental methods ^6-8^. However, overt task behavior can also depend on the maintenance of task-relevant information in awareness ^9^. Accordingly, it is possible that experience sampling can also illuminate the nature of covert processes that are hypothesized to help task performance. Consistent with this possibility, Turnbull and colleagues ^10^ established that when people are off-task, regions involved in external attention, such as regions of the dorsal lateral prefrontal cortex (dLPFC), show reductions in activity, and that in demanding situations, regions of the dLPFC can suppress off-task activity. However, we currently lack an understanding of the patterns of brain activity that support the active maintenance of deliberate task-relevant information covertly in awareness.

To address this gap in our understanding, we re-analyzed a published data set ^11^ to examine whether a pattern of deliberate task-relevant focus is related to specific patterns of neural activity and is linked to better performance on the task at hand. To achieve this goal, we developed a neural state-space based on dimensions of brain function variation that are derived from resting-state connectivity data, and commonly referred to as ‘gradients’ ^12^. These gradients are generated using data-driven techniques and depict axes that differentiate observed function in major brain systems (for a review, see ^13^). We used these gradients to build a 3-d coordinate system that allows us to organize brain maps derived in different ways within a ‘common space’ (see also ^14^). In the current study, we use this common space to examine how patterns of brain activity are related to both 1) covert experiential states that emerge during task processing and that we index via experience sampling and 2) the implementation of different stages of goal-relevant behavior that occur during task completion. The task was a simple sustained attention task in which participants were asked to ignore (frequently presented) non-targets and respond to (infrequent) targets. To index covert experiential states during the task, multidimensional experience sampling (mDES) ^8,15^ was employed, a method that requires individuals to intermittently describe their thoughts by rating several dimensions. This technique has previously been used to identify covert experiential states, including the deliberate maintenance of task-relevant information, as well as patterns of thought that are less related to the here and now, including thoughts with a social episodic focus, and patterns of thought with different modalities (verbal or visual) (e.g., ^10,11,16,17^). This approach is also sensitive to changes in neural function as indexed by functional magnetic resonance imaging (fMRI) ^8,11,14,18-28^.

We used these data to understand whether experience sampling can be used to identify the neural correlates of patterns of thought that are linked to the organization of task-relevant behavior. First, to generate our neural state-space, we selected the first three connectivity gradients ^12^ calculated from resting-state data collected as part of the Human Connectome Project (HCP) ^29^, accounting for approximately 50% of the connectivity variance (see the second panel of Figure 1 for spatial maps representing each gradient). The first connectivity gradient corresponds to a dimension that differentiates sensory-motor cortex from association cortex, the second gradient describes differences between motor and visual cortex, and the third gradient differentiates regions of the default mode network (DMN) from the fronto-parietal network (FPN). The combination of these three gradients forms the 3-d state-space presented in Figure 1. Next, we used these gradients to organize the overt and covert features of cognition derived from our data by projecting the relevant task and experiential maps for each participant into the 3-d space, correlating each map with each gradient dimension. This analytic step resulted in a set of 3 ‘coordinates’ for each task and experiential map for each participant, in which each coordinate describes the location of each map along each gradient dimension. Finally, we used a series of linear mixed models to apply inferential statistics to these coordinates. This analytical pipeline is shown in Figure 1, and see Methods for further details.

**Figure 1.**
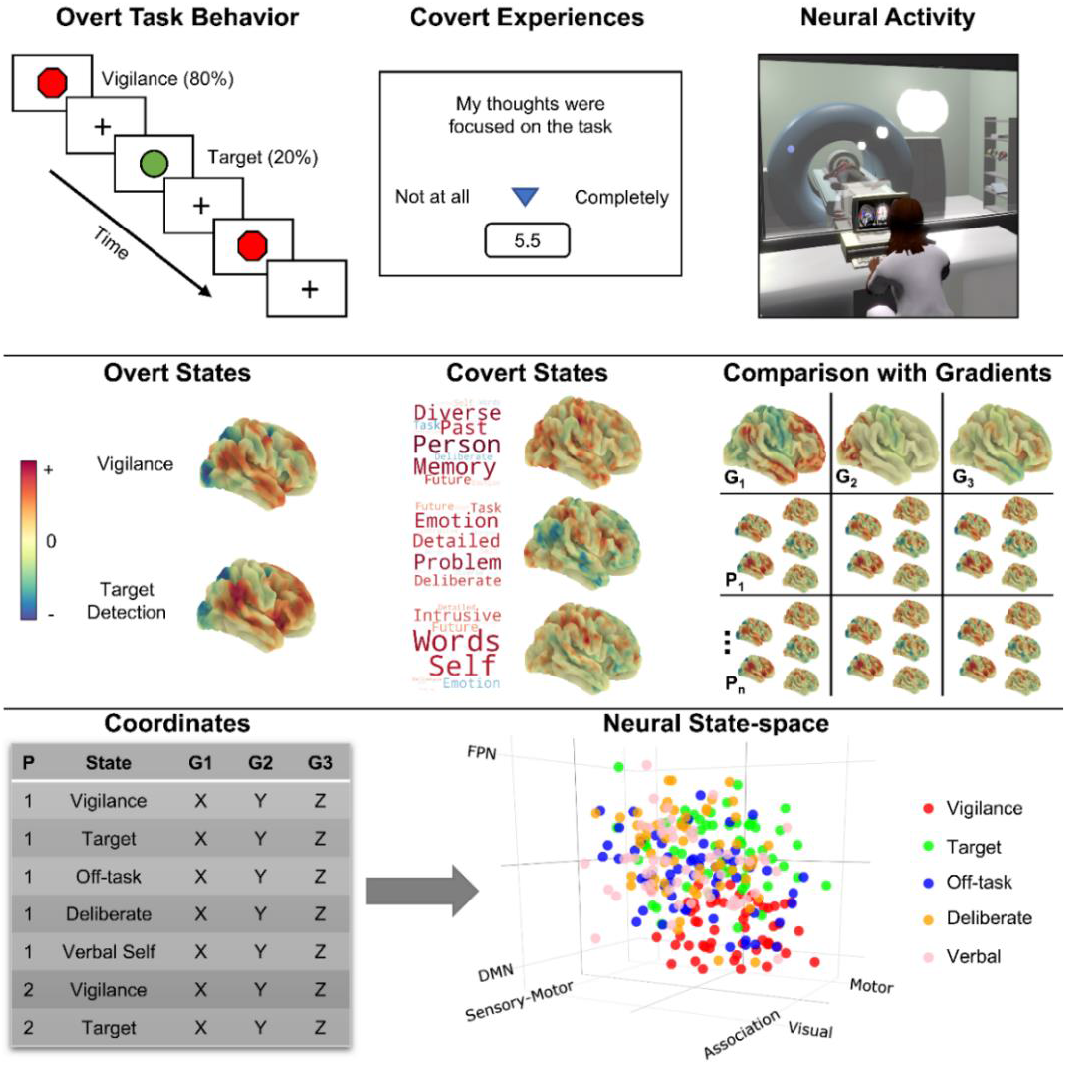
Schematic illustration of the task paradigm and locating the overt task states and covert experiential states in the neural state-space. The top row depicts the task paradigm. Participants completed a sustained attention task while brain activity was measured using fMRI. The task comprised of vigilance periods, in which non-target shapes were presented (80% of trials), and periods of target detection, in which target shapes were presented, requiring a behavioral response (20% of trials). During each run of the task (13 mins x3), 8 experience sampling probes were presented. Each probe asked participants to rate 13 items regarding their covert experience during the task (e.g., level of task focus; see Supplementary Table 5). Principal Components Analysis (PCA) applied to this data identified three thought patterns, represented as word clouds in the second panel (size of word = magnitude of loading and color = direction; warm = positive, cool = negative). Our experimental approach, therefore, allowed us to identify brain correlates to both overt task states (vigilance and target detection) and covert states (as assessed using experience sampling). Brain maps for both overt and covert states (second row) were calculated via the application of the general linear model to each participant using the task time-course, and the time-course for each mDES dimension as explanatory variables (calculated at the first-level using FSL; see Methods). In these brain maps, warmer colors correspond to positive values, while cooler colors correspond to negative value (note, each brain map has a its own color scale). Having identified covert and overt brain states for each individual, we performed pairwise correlations between each of the three connectivity gradients and each individual’s covert and overt brain maps (right-hand side of second row) to produce, for each person and each map, three gradient coordinates, indicating where each individual’s brain maps fall in the 3-d neural state-space (left-hand side of third row). The results of this analysis are shown in the 3-d scatterplot, in which each point represents an individual’s brain maps location in the state-space (N observations = 285). Different colored dots describe different types of brain-cognition relationships.

## Results

### Location of overt task states in the state-space and how these locations predict target detection performance

Our first goal was to determine how the neural state-space differentiated the two features of overt behavior in our task paradigm (vigilance and target detection) and to understand whether their positions in the state-space relate to task performance. To this end, we first calculated the pairwise correlations between spatial brain maps summarizing neural activity during periods of vigilance and target detection with each of the three connectivity gradients, resulting in three sets of coordinates per task map (see Methods). The results of this process can be seen in Figure 2 (panel D), and see Supplementary Figure 1 for the distribution of these coordinates for each task map.

**Figure 2.**
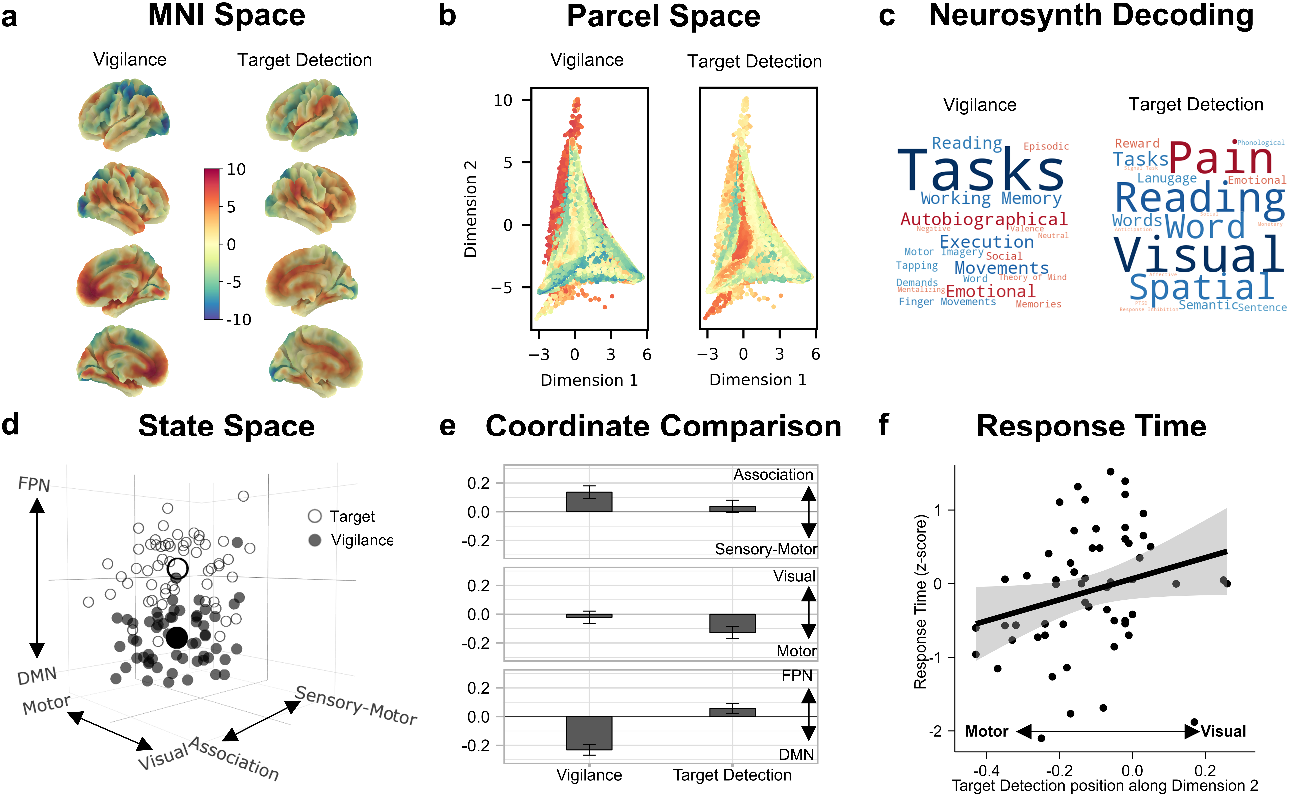
Location of overt task events in the neural state-space and their relationship to task performance. **A)** The (unthresholded) target and vigilance group-level maps plotted on the cortical surface in MNI space (N = 57). These two maps share a common color scale. **B)** A scatter plot of the first two dimensions of the state-space (points = parcels), colored by the fMRI BOLD activity in the target and vigilance brain maps shown in panel A. **C)** Word clouds representing the results from a meta-analysis using NeuroSynth to decode the most likely terms used to describe the pattern of brain activity seen in the target and vigilance maps, where the size of the word represents the magnitude of the relationship, and the color represents the direction (warmer colors = positive relationship, cooler colors = negative relationship). **D)** 3-d scatterplot showing where 1) individual target and vigilance maps fall in the state-space (smaller circles) and 2) the average position of these maps across the sample (larger circles). In this plot, each point represents a whole-brain map for each task condition for each participant (N observations = 114). Open circles represent target maps while closed circles represent vigilance maps. **E)** Bar graph showing the results of the linear mixed models comparing the coordinates of periods of vigilance and moments of target detection along each dimension in the neural state-space. Each bar represents the estimated marginal mean (i.e., predicted value) for each level of ‘task map’ (vigilance and target detection). Error bars represent 95% CIs (N observations = 114). Y-axis labels shown on the right-hand-side indicate the brain systems at the extreme ends of each dimension. **F)** Scatterplot showing the relationship between target detection response time (z-scored seconds; y-axis) and the position of the target map along the motor to visual dimension of the state-space (dimension 2; x-axis). Error bars represents 95% CIs (N observations = 57). X-axis labels indicate extreme ends (motor and vision) of dimension 2.

To understand how these task events are differentiable along the three dimensions of our state-space, we compared the position (i.e., the coordinates) of these maps along each dimension in a series of linear mixed models. We also examined how these positions in the state-space predicted target detection performance during the task in a multiple regression in which the task maps’ coordinates were the explanatory variables and response time was the outcome variable. In addition to traditional significance testing of main effects, bootstrapping (n iterations = 1000) was used to calculate parameter estimates and their associated confidence intervals and p-values to establish the robustness of results emerging from these models (see Methods).

The mixed models comparing the coordinates of each task map along each dimension of the state-space indicated that the position of vigilance and target detection states differed significantly along each dimension [dimension 1: [*F*(1, 56) = 17.01, *P* < .001]; dimension 2: [*F*(1, 56) = 28.30, *P* < .001]; dimension 3: [*F*(1, 56) = 217.86, *P* < .001]]. Along dimension 1—which separates sensory-motor and association cortex—vigilance states, compared to target detection states, were located further towards the association end [bootstrapped *b* = 0.05, 95% CI [0.03, 0.07], *P* < .001]. Along dimension 2—which separates motor and visual systems—target detection states, compared to vigilance states, were located further towards the motor end [bootstrapped *b* = -0.05, 95% CI [-0.07, -0.03], *P* < .001]. Finally, along dimension 3—which separates the default mode and fronto-parietal networks—vigilance states, compared to target detection states, were located further towards the default mode end [bootstrapped *b* = -0.14, 95% CI [-0.16, -0.12], *P* < .001]. Overall, therefore, vigilance states tended to fall towards the association end of dimension 1, the visual end of dimension 2, and the default mode end of dimension 3. In contrast, target detection states tended to fall towards the sensory-motor end of dimension 1, the motor end of dimension 2, and the fronto-parietal end of dimension 3 (see Figure 2E). All bootstrapped parameter estimates for these models are presented in Supplementary Table 1. Finally, the multiple regression indicated that individuals whose target detections states fell further towards the motor end of the motor to visual dimension tended to respond faster to the targets [*F*(1, 47) = 4.47, *P* = .040; bootstrapped *b* = -1.89, 95% CI [-3.35, -0.04], *P* = .044] (see Figure 2F for scatterplot and see Supplementary Table 2 for all bootstrapped parameter estimates).

### Location of covert experiential states in the state-space and how experiential reports predict target detection performance

Having established the position of the overt task states in the state-space, we next made use of this common space to understand the position of the three covert experiential states identified by experience sampling. Principal Components Analysis (PCA) was applied to the mDES data, revealing three components, accounting for 47.41% of the total variance: 1) off-task episodic social cognition, 2) deliberate task focus, and 3) verbal, self-relevant thought (see Methods). These components are represented as word clouds in Figure 3A. We then performed a linear regression, at the individual-level, in which each individuals’ fMRI data during the six seconds prior to the experience-sampling probe was the outcome variable and the individual’s thought score on each of the thought patterns (1-3) was the explanatory variable (see Methods). This produced, for each individual, a spatial map of how their brain activity was associated with their score on each of the three thought patterns identified via PCA. Next, we calculated the pairwise correlations between each of these three ‘experiential’ maps and each of the three connectivity gradients, resulting in three coordinates per experiential map. The results of this process can be seen in Figure 3E, and see Supplementary Figure 1 for the distribution of these coordinates. To understand how these experiential maps are differentiable along the three dimensions of our state-space, we compared the position (i.e., the coordinates) of these maps along each dimension in a series of linear mixed models (see Methods). As before, in addition to traditional significance tests of main effects, we performed bootstrapping (n iterations = 1000) to calculate parameter estimates and their associated confidence intervals and p-values.

**Figure 3.**
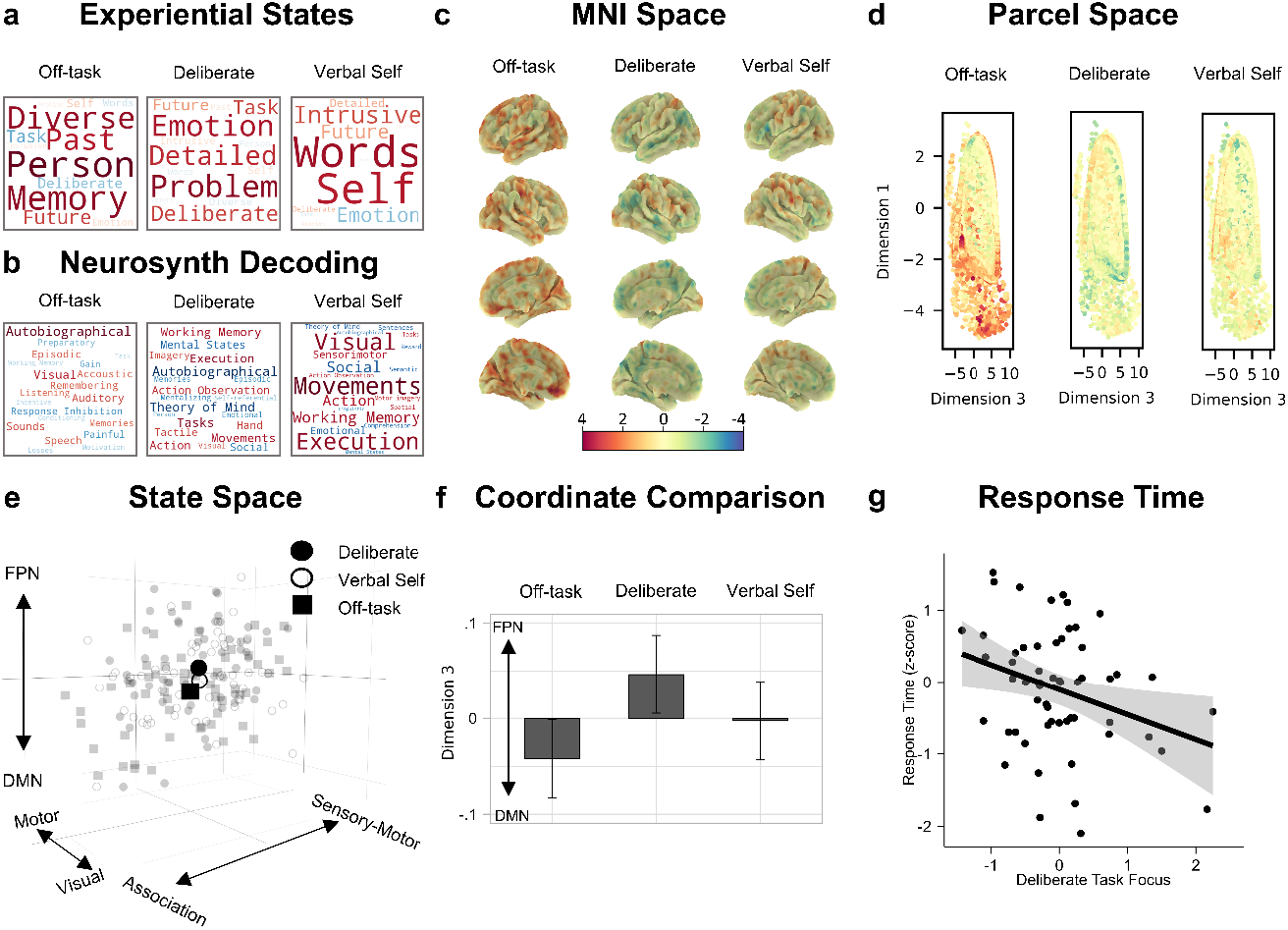
Location of experiential states in the neural state-space and how self-reports of these states predict task performance. **A)** Word clouds representing the three experiential states identified by applying Principal Components Analysis (PCA) to the to the experience sampling data. The first describes patterns of off-task episodic social cognition, the second describes patterns of deliberate task focus, and the third describes patterns of verbal, self-relevant thought. Size of the word represents the magnitude of the relationship, and the color represents the direction (warmer colors = positive relationship, cooler colors = negative relationship). **B)** Word clouds representing the results from a meta-analysis using NeuroSynth to decode the most likely terms used to describe the pattern of brain activity seen in the experiential maps. Size of the word represents the magnitude of the relationship, and the color represents the direction (warmer colors = positive relationship, cooler colors = negative relationship). **C)** The (unthresholded) experiential group-level maps plotted on the cortical surface in MNI space. These three maps share a common color scale. **D)** A scatter plot of the first and third dimensions of the state-space (points = parcels), colored by the fMRI BOLD activity in the experiential brain maps shown in panel C. **E)** 3-d scatterplot showing where 1) individual experiential maps fall in the state-space (smaller points) and 2) the average position of these maps across the sample (larger points). In this plot, each point represents a whole-brain map for each experiential state for each participant (N observations = 171). Squares represent off-task maps, closed circles represent deliberate task focus, and open circles represent verbal, self-relevant thought. **F)** Bar graph showing the results of the linear mixed model comparing coordinates of the experiential states along the DMN to FPN dimension of the state-space. Each bar represents the estimated marginal mean (i.e., predicted value) for each level of ‘experiential map’. Error bars represent 95% CIs (N observations = 171). **G)** Scatterplot showing the relationship between target detection response time (z-scored seconds; y-axis) and the extent to which an individual reported deliberate task focus during the task (x-axis) (N observations = 57).

The mixed models comparing the position of each experiential map along each dimension of the state-space revealed that the position of the experiential maps differed significantly along dimension 3, which separates the default mode and fronto-parietal networks [*F*(2, 165) = 4.75, *P* = .010]. The off-task state fell towards the default mode end of this dimension [bootstrapped *b* = -0.04, 95% CI [-0.07, -0.01], *P* = .008], while the deliberate task focus state fell towards the fronto-parietal end [bootstrapped *b* = 0.04, 95% CI [0.01, 0.08], *P* = .002]. Therefore, our results indicate that covert states of deliberate task focus are associated with patterns of brain activity emphasizing the fronto-parietal network rather than the default mode, whereas, covert off-task states tend to show the opposite pattern. In both cases, the maps were not only different from one another on the default mode – fronto-parietal dimension of the state space, but the distributions of both states fell on average outside the center of this dimension (see Figure 3F). All bootstrapped parameter estimates for these models are presented in Supplementary Table 3.

Finally, we conducted a multiple regression to examine how individual variation in each of the three dimensions of thought was associated with response time. This analysis indicated that patterns of deliberate task focus were associated with faster response times during target detection [*F*(1,50) = 5.51, *P =* .023; bootstrapped *b* = -0.34, 95% CI [-0.60, -0.01], *P* = .046], suggesting that this covert experience may support better task performance (see Figure 3G for scatterplot and see Supplementary Table 4 for all bootstrapped parameter estimates).

## Discussion

Our study set out to examine whether experience sampling can be used to provide insights into covert task-relevant states that are hypothesized to play a role in organizing task behavior. To address this question, we used a simple sustained attention task in which participants detected visually presented targets (to provide an index of overt behavioral states) and experience sampling (to provide indices of covert cognitive states). To allow these different features to be examined in a common analytic space, we used a state-space derived from dimensions of brain function variation ^12^ calculated from the resting state data of the Human Connectome Project ^29^. These dimensions characterize three different types of brain organization: 1) the distinction between sensory-motor and association cortex, 2) the distinction between motor and visual cortex, and 3) the distinction between the default mode and fronto-parietal networks in association cortex. As well as indicating that the default mode network is involved in covert off-task states, experience sampling revealed that the fronto-parietal network is involved in covert states of deliberate focus on the task at hand. Moreover, our data suggest that this pattern of thought was beneficial rather than detrimental to task performance, and the fronto-parietal system was also linked to neural patterns important for the act of target detection. These results, therefore, establish that experience sampling can also describe task-relevant states as well as states that are unconstrained by external input.

As well as establishing the value of experience sampling in understanding task-relevant states, our study provides important insights regarding the mapping between cognition and brain activity. First, our study suggests that association cortex may be generally important when cognition depends on stimulus-independent information processing. We found that patterns of activity within the fronto-parietal and default mode networks distinguished between covert patterns of cognition that are likely to depend, in part, upon information that cannot be inferred directly from immediate sensory input. Contemporary views on the topology of brain organization suggest that regions that make up both the fronto-parietal and default mode networks are located in regions of the cortex that are maximally distant from the systems involved in action and perception ^12^. The location of association cortex at the maximal distance from sensory-motor systems may provide a mechanism through which brain activity in these regions can be more distinct from activity in regions that respond to input describing the immediate environment (for a discussion see: ^30^). Our data, therefore, is consistent with contemporary perspectives on association cortex which suggest that the distance along the cortical mantle from the brain’s input-output systems may be important for helping support patterns of brain activity that are less directly related to information in the immediate environment.

Extrapolating from this perspective, it is possible that different systems within association cortex may serve processes that enable them to help organize cognition and behavior over different time scales. For example, our study suggests that although both the default mode and fronto-parietal networks are important for maintaining covert states, they may do so with a different temporal resolution. Covert states of deliberate thought share organizational features with the patterns of brain activity seen when participants detected a target. This is consistent with a role for this system in maintaining goals relevant to performing the task at hand ^9,31^. In contrast, the default mode network was important for patterns of thought that emphasized mental contents derived from memory and focused on the future or past, instead of the task at hand. These findings are consistent with a role for the default mode network in states like mental time travel ^32-34^, which may be important for organizing behavior over time ^35,36^ and may depend in part on this system’s role in the ability to replay information from memory ^37^. More broadly, this perspective is consistent with the alignment between the default mode network and features of declarative long-term memory such as semantic or episodic memory ^38^.

Although our study demonstrates the value of experience sampling in understanding covert task-relevant states, it nonetheless leaves several questions unanswered. First, since we focused only on one task context, it remains unclear how the current findings generalize to situations with different features. In order to fully understand the role association cortex plays in behavior, it is likely to be particularly important to explore task situations where information from memory is important for guiding actions ^39-41^ and in tasks where performance depends on integration between memory and different types of sensory input (e.g., movie watching or semantic decision making). In the future, therefore, it will be valuable to employ our state-space approach across a wider range of task contexts. This is particularly important given an emerging body of work highlighting the importance of considering the context in which covert experiences emerge when investigating their psychological and neural correlates ^8,16,17,42,43^. Second, since our study focused only on a single scanning session, it will be important for future work to evaluate the consistency and reliability of findings, within and across individuals and over time. For example, future work could use more intensive scanning procedures like those employed in the Midnight Scan Club ^44^ to evaluate the stability of brain state’s locations in the state-space. Third, although we identified a relationship between deliberate thought and better task performance, the significance of this relationship would likely be improved if we could identify situations which maximize the beneficial value of deliberate thought to performance. For example, in a large behavioral sample, Turnbull and colleagues ^45^ demonstrated that off-task thought was more detrimental to performance in a working memory task than in a choice reaction time task. It could be important in the future to understand how variations in deliberate task focus impact on performance across a range of different task conditions to better understand how this thought pattern supports better performance. Finally, although the state-space approach is useful for identifying coarse similarities and differences in whole-brain patterns between states, it may be limited in its ability to identify more fine-grained details. Although this approach is likely to be insufficient to detail modular functions within regions, as our study highlights, it provides a simple way to understand commonalities in brain organization between different types of state and thus is a powerful method to investigate domain general perspectives on how brain organization gives rise to cognition and behavior.

## Materials and Methods

In the following Methods section, the details for the participant information, task paradigm, experience sampling, task procedure, fMRI acquisition and preprocessing, Principal Components Analysis, and FSL-based fMRI analysis are the same as those described in Konu et al. ^11^, with minor rewording in places for clarity (material originally published under a Creative Commons License: <u>CC BY-NC-ND 4.0</u>).

### Participants

One hundred and seven participants took part in this study. Ninety-one participants participated in a behavioral session (67 females; mean age: 23.38 years, standard deviation: 4.53 years, age range: 19–40 years). Sixty-two participants participated in the scanning session (41 females; mean age: 23.29 years; standard deviation: 4.51 years, age range: 18–39 years). After excluding 5 participants, 57 remained for fMRI data analysis (due to technical difficulties or excess movement). Forty-six participants participated in both the behavioral and scanning session. All participants had normal/corrected vision and had no history of psychiatric or neurological illness. All scanning participants were right-handed. This cohort was acquired from the undergraduate and postgraduate student population at the University of York. The study was approved by the local ethics committee at the York Neuroimaging Centre and University of York Psychology Department and all research was performed in accordance with relevant guidelines and regulations. All volunteers provided informed written consent and received monetary compensation or course credit for their participation. These details are the same as those described in Konu et al. ^11^.

### Task paradigm

Participants were instructed to attend to the center of the screen while they were presented with a sequence of ‘non-target’ and ‘target’ stimuli, to which they responded only to target stimuli (mean stimulus presentation duration: 1000 ms). Therefore, this task comprised of ‘vigilance’ periods, in which non-target stimuli were presented, and ‘target detection’ periods, in which target stimuli were presented, requiring a button push. A single run of the task was 13 minutes and contained eight instances of experience sampling probes. For each experience sampling probe, participants rated each experience sampling item once (see the next section for details of the experience sampling technique). Each of the 13 experience sampling items were presented for a maximum of 4 s on the screen—based on the average response time from previous studies—followed by a 500 ms fixation cross. The remainder of the time was allocated to two kinds of experimental trials: target and non-target. In target trials, a green circle was randomly presented (20% of the experiment trials) and participants were required to make a response (a single key or button press). In non-target trials, a red octagon was presented (80% of the experiment trials) and no behavioral response was required. An experimental trial was fixed at 3000 ms. The inter-stimulus-intervals (ISI) consisted of a fixation cross and was jittered (1500–2500 ms). The stimulus was presented on screen for 500–1500 ms until a response was made. Once a response was captured, a fixation cross appeared on the screen for the remaining time. This task was designed to require minimal cognitive demand since these conditions (i.e., long periods of vigilance, interleaved with simple target detection) facilitate the occurrence of self-generated thought at a level that is comparable to rest ^46^. The task paradigm is presented schematically in the top panel of Figure 1. In the scanner, participants completed three runs of the task, whereas, in the behavioral session they completed one run of the task. Written instructions were presented at the start of each run. These details are the same as those described in Konu et al. ^11^.

### Multidimensional Experience Sampling (mDES)

Participants’ ongoing thought throughout the task was measured using a technique known as multidimensional experience sampling (mDES) ^15^. When an mDES probe occurred, participants were first asked how much their thoughts were focused on the task, followed by 12 randomly shuffled items about the content and form of their thoughts (see Supplementary Table 5 for all mDES items). All items were rated on a 1-10 continuous scale (see top panel of Figure 1 for an illustration). Within one run of the task, participants completed 8 sets of mDES probes, yielding a total of 8 probes per individual in the behavioral session and 24 probes per individual in the scanning session. In the scanning session, two participants had one run dropped due to technical issues, leaving them with 16 probes overall. These details are the same as those described in Konu et al. ^11^.

### Procedure

In the behavioral session, participants completed a single 13-min run of the task with mDES. In the scanning session, participants completed three, 13-min functional runs of the task with mDES while undergoing fMRI. The scanner session took around 1 h and 15 min, of which the task took ∼45 min, and this was separated into three blocks. These details are the same as those described in Konu et al. ^11^.

### fMRI acquisition

All MRI scanning was carried out at the York Neuroimaging Centre. Structural and functional scans were acquired using a Siemens Prisma 3T MRI Scanner with a 64-channel phased-array head coil. Structural data were acquired using a T1-weighted (MPRAGE) whole-brain scan (TR = 2300 ms, TE = 2.26 ms, flip angle = 8°, matrix size = 256 × 256, 176 slices, voxel size = 1 × 1 × 1 mm). Functional data were collected using a gradient-echo EPI sequence with 54 bottom-up interleaved axial slices (TR = 3000 ms, TE = 30 ms, flip-angle = 80°, matrix size = 80 × 80, voxel size = 3 × 3 × 3 mm, 267 vol) covering the whole brain. These details are the same as those described in Konu et al. ^11^.

### fMRI data pre-processing

Functional and structural data were pre-processed and analyzed using FMRIB’s Software Library (FSL, version 5.0.1, http://fsl.fmrib.ox.ac.uk/fsl/fslwiki/FEAT/). Individual T1-weighted structural images were extracted using BET (Brain Extraction Tool). Functional data were pre-processed and analyzed using the FMRI Expert analysis Tool (FEAT). Individual participant analysis involved motion correction using MCFLIRT and slice-timing correction using Fourier space time-series phase-shifting. In the current study, to control for individual’s movement during the scanning period in our inferential analyses, we also calculated each individual’s mean movement across all three runs using the MCFLIRT output to include as a nuisance regressor in subsequent analyses (prefiltered_func_data_mcf_abs_mean.rms). After co-registration to the structural images, individual functional images were linearly registered to the MNI-152 template using FMRIB’s Linear Image Registration Tool (FLIRT). Registration from high resolution structural to standard space was then further refined using FNIRT nonlinear registration. Functional images were spatially smoothed using a Gaussian kernel of FWHM 6 mm, underwent grand-mean intensity normalization of the entire four-dimensional dataset by a single multiplicative factor, and had high pass temporal filtering (Gaussian-weighted least-squares straight line fitting, with sigma = 50s). These details are the same as those described in Konu et al. ^11^.

### Principal Component Analysis

Analysis of the mDES data was carried out in SPSS (Version 25, 2019). Principal Component Analysis (PCA) was applied to the scores from the 13 experience-sampling items comprising the probes for each participant. PCA was applied at the trial level in the same manner as in our prior studies e.g., ^15-17,23,47,48,49^. Specifically, we concatenated the responses of each participant for each trial into a single matrix and employed a PCA with varimax rotation. We performed this analysis separately for each session (behavioral and scanning) in order to examine the similarity in the solutions produced across each situation (see Supplementary Figure 2). These details are the same as those described in Konu et al. ^11^. In the current study, we calculated each individual’s mean score, across runs, for each PCA component identified for inclusion in inferential analyses. Intraclass correlation (ICC) analyses indicated moderate consistency of component 1 (0.68), and good consistency of component 2 (0.77) and component 3 (0.75) across the three runs.

### fMRI analysis

#### Creating brain maps for covert experiential states and overt task states

Task-based analyses were carried out using FSL. A model was set up including 6 explanatory variables (EVs). EVs 1 and 2 modeled ‘vigilance’ and ‘target detection’ periods. EV 3 modeled activity 6s prior to each mDES probe. Finally, EVs 4, 5, and 6 modeled the 3 thought components identified through PCA, with a time period of 6s prior to the mDES probes and the scores for the relevant component as a parametric regressor. EVs were mean-centered within each run and no thresholding was applied to the EVs. Standard and extended motion parameters were included as confounds. This was convolved with a hemodynamic response function using FSL’s gamma function. We chose to use the same 6s interval as used in Turnbull et al. ^23^. Contrasts were included to assess brain activity that related to each of the two task events (vigilance and target detection) and that related to each component of thought during the 6s period prior to the probe. The three runs were included in a fixed-level analysis to average across the activity within an individual. The averaged-run individual-level unthresholded (z-stat) contrast maps were used in the state-space analyses described below. Group-level analyses followed best practice ^50^. Specifically, we used FLAME, as implemented by FSL. Figure 2A and Figure 3C show the unthresholded group-level maps in MNI space while Figure 2B and Figure 3D show scatterplots representing how the BOLD activity in the maps’ parcels is distributed along the state-space dimensions. The unthresholded group-level maps were used in the NeuroSynth analysis described in the next section. These details are the same as those described in Konu et al. ^11^.

#### Neurosynth decoding of covert experiential and overt task spatial maps

We used Neurosynth’s online meta-analytical decoder ^51^ to identify terms most commonly associated with our (unthresholded) group-level neural maps in the available literature (5 maps: vigilance, target detection, off-task thought, deliberate thought, and verbal, self-relevant thought). When provided with an unthresholded whole-brain map, the Neurosynth decoder performs a reverse inference analysis to identify the terms describing cognitive and psychological functions that are most strongly associated with the patterns of neural activation shown in the map. Specifically, the decoder compares the patterns of activation in the input map to patterns of activation in the studies in the Neurosynth database, resulting in a list of terms that are most likely to be positively or negatively associated with the patterns of activation in the input map. To visualize the results of this analysis as word clouds, we selected the top 10 positive and top 10 negative cognitive and psychological terms associated with each map, retaining only the first term in instances of duplicates (e.g., ‘autobiographical’ and ‘autobiographical memory’) and excluding terms related to anatomy instead of function (e.g., ‘occipital’). These word clouds are shown in Figure 2C (overt states) & Figure 3B (covert states).

#### Connectivity gradients used to construct the 3-d neural state-space

The three connectivity gradients used in the current study were generated by Margulies et al. ^12^ and are openly available via Neurovault: https://identifiers.org/neurovault.collection:1598. These gradients were generated by applying a non-linear dimension reduction technique (diffusion embedding) to the averaged functional connectivity matrix of the Human Connectome Project (HCP) data ^29^. These gradients explain whole-brain connectivity variance in descending order, such that the first gradient explains the most variance in the whole-brain connectivity data, the second explains the second most variance, and so on. Along each gradient, brain regions with similar connectivity profiles (to the rest of the brain) fall close together, and have similar ‘gradient values’, while regions with more distinct connectivity profiles fall further apart, and have more dissimilar ‘gradient values’ ^13^. This analysis, therefore, results in a spatial map for each gradient identified in which each parcel contains a ‘gradient value’. Prior studies have highlighted that the first three gradients relate to important features of cognition ^14,40,52^. We use these three gradients to construct the 3-d neural state-space (see below). The first gradient describes the difference between sensory-motor regions and association cortex. The second gradient separates motor and visual systems. Finally, the third gradient describes the difference between the DMN and the FPN (see second panel of Figure 1).

#### Locating overt task states and covert experiential states in the neural state-space

To locate overt task states and covert experiential states in the neural state-space, we calculated the pairwise spatial correlations (Pearson) between each individual’s covert and overt brain maps and each of the first three connectivity gradients described in Margulies et al. ^12^. Therefore, for each individual, this resulted in three correlation values for each brain map, indicating where that brain map falls along each dimension of the neural state-space. These correlation values act as ‘coordinates’ in the 3-d neural state-space (see Figure 1). Finally, we Fisher-z transformed the correlation values before using them in inferential analyses.

#### Linear Mixed Models

Linear Mixed Models (LMMs) were fitted by restricted maximum-likelihood estimation in R [4.1.1 ^53^] using the lme4 package [1.1.31 ^54^]. We used the lmerTest package [3.1.3 ^55^] to obtain *P* values for the F-tests returned by the lme4 package. For each set of models, the alpha level for each F-statistic was set based on 0.05 divided by the number of models (i.e., Bonferroni-corrected alpha level). Degrees of freedom were calculated using Satterthwaite approximation and for F-tests, type 3 sum of squares was used. Contrasts were set to “contr.sum,” meaning that the intercept of each model corresponds to the grand mean of all conditions and that when a factor has two levels, the parameter estimate is equal to half of the difference between the two levels ^56^. Estimated marginal means (shown in Figure 2E and 3F) were calculated using the emmeans package [1.8.3 ^57^]. Across all models, to account for multiple observations per participant, ‘participant’ was included as a random intercept. Finally, to establish the robustness of our results, we used the easystats package [0.6.0 ^58^] to obtain bootstrapped parameter estimates (n iterations = 1000) and their associated confidence intervals and *P* values.

#### Comparing the location of overt task states in the neural state-space

We ran three LMMs—one with each dimension coordinate as the outcome variable and “task map” as the explanatory variable (two levels: vigilance and target detection). Age, gender, and mean movement were included as nuisance covariates. In total, 57 participants were included in these models. These models allowed us to investigate how the location of the two overt task states (vigilance and target detection) differed along each dimension of the neural state-space.

Example model formula: lmer(Dimension Coordinate X ∼ Task Map + Age + Gender + Mean Movement + (1|Participant))

#### Comparing the location of covert experiential states in the neural state-space

We ran three LMMs—one with each dimension coordinate as the outcome variable and “experiential map” as the explanatory variable (three levels: off-task thought, deliberate thought, and verbal, self-relevant thought). Age, gender, and mean movement were included as nuisance covariates. In total, 57 participants were included in these models. These models allowed us to investigate how the location of the three covert experiential states (off-task, deliberate, and verbal, self-relevant thought) differed along each dimension of the neural state-space.

Example model formula: lmer(Dimension Coordinate X ∼ Experiential Map + Age + Gender + Mean Movement + (1|Participant))

#### Multiple Regressions

Multiple regressions were fitted by ordinal least squares (OLS) in R [4.1.1 ^53^] using the stats package. We used the rstatix package [0.7 ^59^] to obtain F-test statistics and associated p-values. For F-tests, type 3 sum of squares was chosen and contrasts were set to “contr.sum,” meaning that the intercept of each model corresponds to the grand mean of all conditions and that when a factor has two levels, the parameter estimate is equal to half of the difference between the two levels ^56^. To establish the robustness of our results, we used the easystats package [0.6.0 ^58^] to obtain bootstrapped parameter estimates (n iterations = 1000) and their associated confidence intervals and *P* values. Across all models, age, gender, and mean movement were included as nuisance covariates. Finally, in these reaction time models, cases exhibiting a z-scored response time greater than 2.5 were considered outliers, and the z-scores of these outliers were set to zero to mitigate their influence on the results. Using this approach, two cases were considered outliers.

#### Examining how the location of overt task states in the neural state-space predict target detection reaction time

We ran a multiple regression in which response time (z-scored) was the outcome variable and the three coordinates for each of the two overt task brain maps (vigilance and target detection) were the explanatory variables (6 in total). In total, 57 participants were included in these models. This regression allowed us to investigate whether there is a correspondence between the location of overt task states within the neural state-space and target detection performance.

Example model formula: lm(Z-scored Response Time ∼ Target Detection Coordinate along Dimension 1 + Target Detection Coordinate along Dimension 2 + Target Detection Coordinate along Dimension 3 + Vigilance Coordinate along Dimension 1 + Vigilance Coordinate along Dimension 2 + Vigilance Coordinate along Dimension 3 + Age + Gender + Mean Movement)

#### Examining how experiential reports of covert states predict target detection reaction time

We ran a multiple regression in which response time (z-scored) was the outcome variable and individual’s mean scores for each of the three thought patterns (off-task thought, deliberate thought, and verbal, self-relevant thought) were the explanatory variables. In total, 57 participants were included in these models. This model allowed us to investigate whether there is a correspondence between individual’s covert experiences and their target detection performance.

Example model formula: lm(Z-scored Response Time ∼ Off-task Thought + Deliberate Thought + Verbal Self-relevant Thought + Age + Gender + Mean Movement)

## Data Availability

Ethical approval conditions and European Research Council (ERC) grant stipulations do not permit the public sharing of raw data. However, anonymized Multidimensional Experience Sampling data and Gradient coordinates are publicly available via Mendeley: https://doi.org/10.17632/mx76fvdm3v.1. In addition, all group-level unthresholded brain maps presented in the figures are available via NeuroVault: https://identifiers.org/neurovault.collection:13520. The code for the task paradigm is publicly available at: https://vcs.ynic.york.ac.uk/hw1012/go_nogo_experience_sampling/tree/master/. All code used in the current analysis and preparation of figures is publicly available via GitHub at: https://github.com/Bronte-Mckeown/mDES_States.

## Acknowledgments

Chat GPT was used to improve the explanation of the NeuroSynth decoder in Methods. This project was supported by the European Research Council Consolidator Grant awarded to J.S. (WANDERINGMINDS–646927).

## Author Contributions

DK, HTW, EJ, & JS designed task paradigm; DK collected the data; MZ, TK, TX, & DM contributed analytical tools; BM, WHS, AT, RL, BB, DM, JW & JS conceived and designed the analysis; BM analyzed data; BM prepared visualization; BM & JS writing-original draft; BM, WHS, MZ, AT, DK, TK, HTW, RL, TX, SH, BB, DM, EJ, JW & JS writing-review and editing; JS acquired funding.

## Additional Information

The authors declare no competing interest.

