## Supplementary materials for "Experience sampling reveals the role that covert goal states play in task-relevant behavior"

**This file includes:**

Supplementary Figures 1 to 2
Supplementary Tables 1 to 5

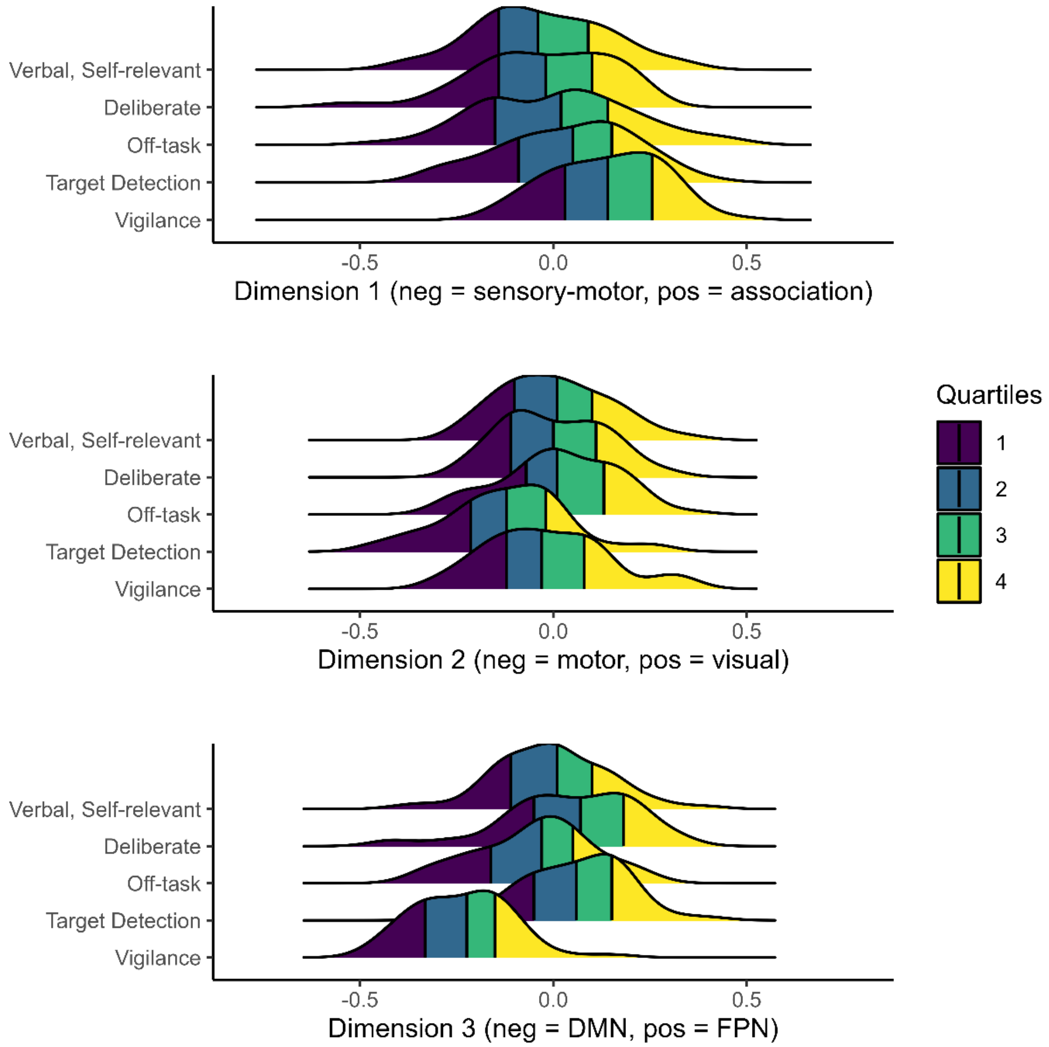


**Supplementary Figure 1.** Density ridge plots showing the distribution of each task and experiential states' coordinates along each dimension of the neural state-space. The y-axis shows each of the five brain states examined in the current study (3 covert states, 2 overt states), and the x-axis shows the coordinate values along each dimension. The color indicates 25% Quartiles. N observations per brain state = 57. In this plot, 'pos' = positive values, 'neg' = negative values, DMN = default mode network, and FPN = fronto-parietal network.

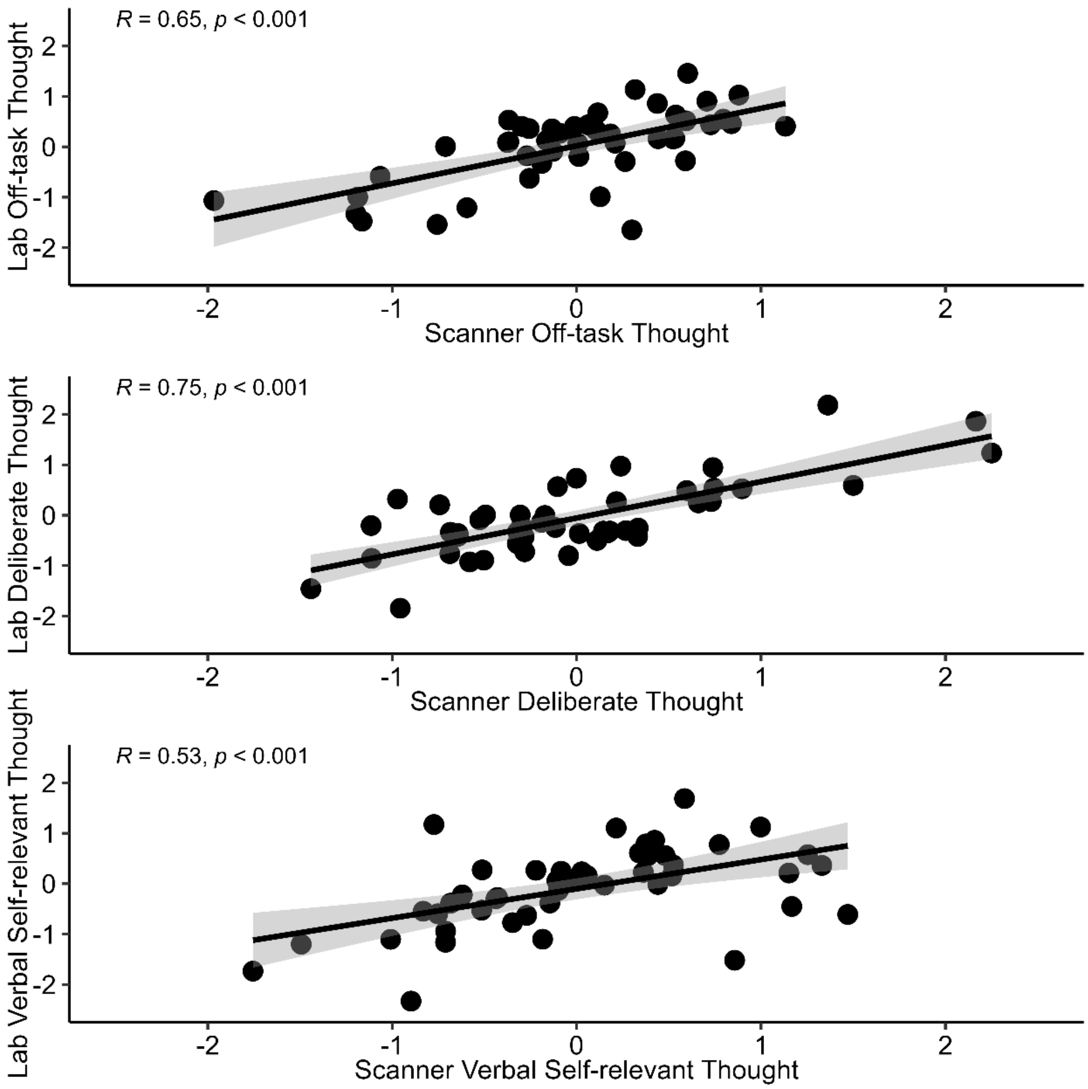


**Supplementary Figure 2.** Scatterplots showing the correlations between average PCA scores derived from Principal Components Analysis (PCA) applied to the experience sampling data collected in 1) the lab (y-axis) and 2) the scanner (x-axis). Error bars represent 95% Confidence intervals. N observations = 46.

**Supplementary Table 1.** Bootstrapped estimates (n iterations = 1000) for linear mixed models comparing the location of overt task states (vigilance & target detection) in the neural state-space.

| | Dimension 1 | | | | *Dimension 2* | | | | Dimension 3 | | | |
|---|---|---|---|---|---|---|---|---|---|---|---|---|
| *Parameter* | *b* | *Lower 95 CI* | *Upper 95 CI* | *p* | *b* | *Lower 95 CI* | *Upper 95 CI* | *p* | *b* | *Lower 95 CI* | *Upper 95 CI* | *p* |
| (Intercept) | 0.03 | -0.16 | 0.21 | .726 | -0.04 | -0.24 | 0.15 | .618 | -0.08 | -0.23 | 0.07 | .296 |
| Vigilance | 0.05 | 0.03 | 0.07 | **<.001** | 0.05 | 0.03 | 0.07 | **<.001** | -0.14 | -0.16 | -0.12 | **<.001** |
| Female | 0.00 | -0.04 | 0.04 | .922 | 0.02 | -0.02 | 0.06 | .290 | -0.02 | -0.05 | 0.01 | .288 |
| Age | 0.00 | -0.00 | 0.01 | .504 | -0.00 | -0.01 | 0.01 | .664 | -0.00 | -0.01 | 0.01 | .866 |
| Mean Movement | -0.01 | -0.11 | 0.10 | .840 | 0.03 | -0.09 | 0.13 | .666 | 0.01 | -0.08 | 0.10 | .802 |

*Note. P* < .05 highlighted in bold. Summed contrasts were used meaning that the intercept reflects the grand mean of all conditions for each model, each factor level estimate reflects the difference between the factor level and the intercept and that when a factor has two levels, the parameter estimate is equal to half of the difference between the two levels.

**Supplementary Table 2.** Bootstrapped estimates (n iterations = 1000) for the multiple regression examining how the location of overt task states (vigilance and target detection) in the neural state-space predict target detection reaction time.

| *Parameter* | *b* | *Lower 95 CI* | *Upper 95 CI* | *p* |
|---|---|---|---|---|
| (Intercept) | 0.28 | -1.42 | 1.92 | .744 |
| Vigilance along dimension 1 | -1.14 | -3.51 | 1.01 | .278 |
| Vigilance along dimension 2 | -0.88 | -2.92 | 1.13 | .434 |
| Vigilance along dimension 3 | -0.22 | -2.74 | 2.32 | .850 |
| Target detection along dimension 1 | 0.22 | -1.37 | 2.24 | .808 |
| Target detection along dimension 2 | 1.89 | 0.04 | 3.35 | **.044** |
| Target detection along dimension 3 | -0.77 | -2.76 | 1.03 | .424 |
| Age | -0.00 | -0.06 | 0.05 | .932 |
| Female | 0.23 | -0.02 | 0.49 | .066 |
| Mean Movement | 0.03 | -0.83 | 0.48 | .888 |

*Note*. *P* < .05 highlighted in bold. Reaction time was z-scored. Summed contrasts were used meaning that the intercept reflects the grand mean of all conditions for each model and each factor level estimate reflects the difference between the factor level and the intercept.

**Supplementary Table 3.** Bootstrapped estimates (n iterations = 1000) for linear mixed models comparing the location of covert experiential states (off-task episodic social cognition, deliberate task focus, and verbal, self-relevant thought) in the neural state-space.

| | Dimension 1 | | | | Dimension 2 | | | | Dimension 3 | | | |
|---|---|---|---|---|---|---|---|---|---|---|---|---|
| *Parameter* | *b* | *Lower 95 CI* | *Upper 95 CI* | *p* | *b* | *Lower 95 CI* | *Upper 95 CI* | *p* | *b* | *Lower 95 CI* | *Upper 95 CI* | *p* |
| (Intercept) | 0.04 | -0.12 | 0.18 | .620 | -0.01 | -0.12 | 0.10 | .858 | -0.03 | -0.15 | 0.10 | .676 |
| Off-task | 0.03 | -0.01 | 0.07 | .194 | 0.01 | -0.02 | 0.04 | .588 | -0.04 | -0.07 | -0.01 | **.008** |
| Deliberate | -0.01 | -0.05 | 0.03 | .534 | -0.00 | -0.04 | 0.02 | .810 | 0.04 | 0.01 | 0.08 | **.002** |
| Female | 0.00 | -0.02 | 0.03 | .848 | -0.01 | -0.03 | 0.02 | .562 | -0.01 | -0.04 | 0.02 | .512 |
| Age | -0.00 | -0.01 | 0.01 | .692 | 0.00 | -0.00 | 0.01 | .544 | 0.00 | -0.00 | 0.01 | .478 |
| Mean Movement | -0.07 | -0.16 | 0.00 | .066 | -0.03 | -0.10 | 0.03 | .318 | -0.04 | -0.11 | 0.03 | .306 |

*Note. P* < .05 highlighted in bold. Summed contrasts were used meaning that the intercept reflects the grand mean of all conditions for each model and each factor level estimate reflects the difference between the factor level and the intercept.

**Supplementary Table 4.** Bootstrapped estimates (n iterations = 1000) for the multiple regression examining how experiential reports of covert states predict target detection reaction time.

| *Parameter* | *b* | *Lower 95 CI* | *Upper 95 CI* | *p* |
|---|---|---|---|---|
| (Intercept) | -0.36 | -1.60 | 0.75 | .556 |
| Off-task | 0.17 | -0.25 | 0.50 | .344 |
| Deliberate | -0.34 | -0.60 | -0.01 | **.046** |
| Verbal Self | 0.01 | -0.33 | 0.29 | .930 |
| Age | 0.01 | -0.03 | 0.06 | .696 |
| Female | 0.21 | -0.02 | 0.44 | .070 |
| Mean Movement | -0.09 | -1.09 | 0.39 | .664 |

*P* < .05 highlighted in bold. Reaction time was z-scored. Summed contrasts were used meaning that the intercept reflects the grand mean of all conditions for each model and each factor level estimate reflects the difference between the factor level and the intercept.

**Supplementary Table 5.** Multidimensional Experience Sampling (mDES) items used in the current study to examine the contents and form of individuals' ongoing thoughts during the task.

| Dimension | Statement | Scale low | Scale high |
|---|---|---|---|
| Task | My thoughts were focused on the task I was performing: | Not at all | Completely |
| Future | My thoughts involved future events: | Not at all | Completely |
| Past | My thoughts involved past events: | Not at all | Completely |
| Self | My thoughts involved myself: | Not at all | Completely |
| Person | My thoughts involved other people: | Not at all | Completely |
| Emotion | The emotion of my thoughts was: | Negative | Positive |
| Modality | My thoughts were in the form of: | Images | Words |
| Detail | My thoughts were detailed and specific: | Not at all | Completely |
| Deliberate | My thoughts were: | Spontaneous | Deliberate |
| Problem | I was thinking about solutions to problems (or goals): | Not at all | Completely |
| Diverse | My thoughts were: | One topic | Many topics |
| Intrusive | My thoughts were intrusive: | Not at all | Completely |
| Source | My thoughts were linked to information from: | Environment | Memory |
